# Variants in glycine decarboxylase activate mechanisms of mitochondrial energy metabolism in the brain

**DOI:** 10.1101/2025.07.12.664515

**Authors:** Alejandro Lopez-Ramirez, Andrew J Worth, Ziyi (Lindsay) Wang, Saliha Yilmaz, Sebastian Hayes, Joseph Farris, Md Suhail Alam, Prasad Padmanabhan, Kasturi Haldar

## Abstract

Brain energy metabolism is produced from glucose by mitochondrial oxidative phosphorylation. Variants in the mitochondrial enzyme glycine decarboxylase (GLDC) cause a rare neurological disease, non-ketotic hyperglycinemia (NKH), with expected hallmarks of brain glycine elevation and responsiveness to folate deficiency that are equivalent to the severity of *Gldc* mutations. We remarkably find that brains of young- attenuated mutant mice with a 1.5-fold increase in glycine are reduced > 5-fold in GLDC, show a decline in both the mitochondrial lipoyl-transfer protein GCSH and lipoylation of the pyruvate dehydrogenase (PDH) complex, as well as concomitant rise in signatures of astrocyte mitochondrial β-oxidation of fatty acids and activation of neuronal PDH. Our findings suggest a novel GLDC mechanism of remodeling mitochondrial energy systems throughout the brain, established early in and sustained throughout post-natal NKH disease.

## Introduction

The brain requires high levels of energy, utilizing a fifth of the total required by the body^1^. 95% of the brain’s energy comes from glucose^1–4^. Oxidative phosphorylation of glucose via the mitochondria produces acetyl co-enzyme A (acetyl-CoA) the key precursor for energy, whose production and levels are regulated by the pyruvate dehydrogenase (PDH) complex. However, glucose levels within the brain are sufficient to provide energy for only a few minutes in neurons. Therefore, energy challenges create the need for alternate fuels sourced from outside. Neurons also receive energy from primary glial cells called astrocytes, which depend on glycolysis for their intrinsic metabolic functions^5–8^. Lactate from astrocytes is an important energy source for neurons ^2,9,10^. Unlike neurons, astrocytes have the capacity to mitochondrially support β-oxidation of fatty acids as an alternate energy source, mitigation of which is known to impair cognition^11–16^. Brain glucose metabolism and mitochondrial oxidative phosphorylation weaken in neurodegenerative disease and cumulative evidence suggests that they are important for both neuronal and glial functions ^17^.

Nonketotic Hyperglycinemia (NKH) is a rare neurometabolic disorder caused by a defect in the Glycine Cleavage System (GCS), which is a major pathway for glycine catabolism. The global prevalence of NKH is estimated at 1 in 76,000^18^. Approximately 80% of NKH is caused by a defect in glycine decarboxylase (GLDC). NKH is a complex, poorly understood disease with a wide range of clinical severity and over 50 different symptoms ^19,20^. Patients with attenuated mutations and disease are affected by intellectual disability and ADHD-like characteristics^21^, while those with severe disease often display intractable seizures, loss of consciousness, and severe hypotonia ascribed to high levels of glycine in the brain^18,20,22,23^. However, in murine models, elevation of brain glycine alone is insufficient to induce severe neurological disease ^24,25^. Deficiency in GLDC also disrupts formation of 5, 10-methyl tetrahydrofolate (5,10-meTHF), a key intermediate in folate metabolism that gives rise to fatal, pre-natal hydrocephalus^26^. But whether there are direct effects on the mitochondrial PDH complex that is critical for the support of brain energy metabolism is not known.

Since GLDC has over 200 variants, in prior studies, we developed large scale genotype-phenotype studies of human mutations and quantitative tools to comprehensively predict and separate severe versus attenuated mutations based on their associated clinical disease status^27^. We then successfully validated these predictions using CRISPR Cas9 to engineer mice with attenuated and highly pathogenic mutations and develop humanized mouse models of attenuated and severe NKH disease^28^. The neuro-metabolic range captured between the attenuated and severe mice suggested capacity to assess phenotypic determinants of mutant mice at either end of the disease spectrum.

Although GLDC is a mitochondrial enzyme, defect in which has neurological consequences, its direct effect on mechanisms of brain energy metabolism has never been evaluated either clinically or in model systems. GLDC is reported to be present across all major regions of the brain^29^ . Both current and historic evidence concur that it is highly expressed in astrocytes ^30,31^, which may comprise up to 40% of all cells in the brain. Expression of GLDC protein in microglia (which account for ∼10% of cells in the brain) is not supported by published literature. One recent study suggests the presence of low levels of GLDC in subtypes of neurons ^32^. In the mouse brain, GLDC is principally found in astrocytes, depleted in microglia, and present at extremely low levels in neurons and oligodendrocytes^33^. We have recently shown in a humanized NKH mouse model expressing a prevalent *Gldc* attenuated disease mutation^28^, that astrocytes (but not neurons, microglia or oligodendrocytes) are reduced between one to five months of age^34^. This strongly supports primary action of *Gldc* deficiency is directly detrimental to astrocytes. Hence, NKH mice may provide model systems to separate how mitochondrial mechanisms of energy metabolism of astrocytes and neurons are regulated by GLDC, all of which remain undiscovered.

## Materials and Methods

### Animals

CRISPR-Cas9 gene editing was used to introduce the attenuated (A394V, referred to as *Gldc^ATT/ATT^*) mutation in the C57BL6 strain ^28^. A restriction enzyme site (GTGAAC) for Hpy166II was created in the mutant allele. Tail biopsies collected and genotyping by PCR- amplifying a 753 bp DNA fragment of *Gldc* flanking the mutation site using the following primers (forward: 5’-GTTGCATTTCCGTTTCTGGCT-3’ and reverse: 5’-ACTGCCCTCTTACTTGACCATT-3’), digesting the amplified product with Hpy166II and separating fragments by agarose gel electrophoresis.

Severe mutations (G561R, S562F) mutations in the *Gldc* gene were introduced by CRISPR-Cas9 editing into C57BL/6 mice to create the *GldcSEV/SEV* strain ^28^. To create a restriction site for BsrG1, two silent nucleotide substitutions (TGCACC in wild type changed to TGTACA in the mutant), were introduced downstream of the knock-in site. Genotyping was carried out as follows. A 728 bp DNA fragment of *Gldc* was PCR amplified using primers (forward: 5’-TGCTGTGCTGGGGAGAATTT-3’ and reverse: 5’- TGAACACAGCTACACTCAGCTT-3’), the amplified production was digested with BsrG1 and fragments were separated by gel electrophoresis. As previously reported^28^ in the breeding of severe mutants, after 48h of pairing, dams were separated and sodium formate (30mg/ml) was dissolved in Sucralose (Portland, ME, USA) and provided in the drinking water: fomate-supplementation was discontinued once pups were born.

### Mouse brain proteome and metabolite analyses

Mouse brains were isolated and flash frozen and stored at -80°C.

### Brain Proteome Analyses

Tissue sample preparation, processing, and analysis for proteome analysis were undertaken as previously described^35^. **Quality Control (QC) and Normalization**. Proteomic data were subjected to rigorous pre- processing steps to ensure data quality and reliability. Initially, features (proteins) with zero or missing values (NA) across all groups were excluded from the analysis. Subsequently, variance stabilizing normalization (vsn)^36^ was applied to the dataset to correct for any systematic biases and to stabilize the variance across the measurements. Samples exhibiting low correlation within their respective groups were identified and removed from the dataset if their correlation coefficient fell below a threshold of 0.8. In this study, one sample was removed based on this criterion. Additionally, features with low repeatability, characterized by a coefficient of variation greater than 0.3, were excluded to maintain data robustness. Missing values in the dataset were imputed using the mean value of all available samples to facilitate subsequent analyses. After quality control steps, the brain proteomics dataset consisted of 5975 features across 65 samples, reduced from an initial 6254 features before QC.

### Statistical Analysis of Differential Protein Expression and clustering

All statistical analyses were performed using R language in the RStudio environment. We conducted age-matched two-sample t-tests for each proteinin analyses that were stratified by phenotype and age group. Initially, nominal p-values less than 0.01 were considered significant and the hits were subsequently subjected to multiplicity correction using the Benjamini-Hochberg method b.Tissue metabolite analysis by LC-MS Pulverized frozen tissue was spiked with 500 ng [^13^C2^15^N]-glycine and [^13^C_3_ ^15^N]-serine (Sigma) and extracted with ice cold 80/20 MeOH/water (40 µL/mg tissue). For absolute quantification of L/D-serine and glycine, a stardard curve of each was run against the same 500 ng internal standard. Samples were spun at 10,000xg for 10 min and supernatants transferred and dried prior to reconstitution in 50 µL water for LC-MS analysis (10 µL injection volume).

Positive and negative mode metabolite analyses were performed with reversed-phase ion-pairing liquid chromatography mass spectrometry (LC-MS) on a Thermo Vanquish Flex pump coupled to a QExactive orbitrap mass spectrometer using electrospray ionzation (Thermo Fisher Scientific, San Jose, CA). Chromatography for negative mode ionization, the stationary phase was an ACQUITY UPLC HSS T3 (1.8 μm 2.1x150 mm) column. LC separation was achieved with a gradient elution of solvent A (97/3 H2O/methanol with 10 mM tributylamine, 15 mM acetic acid at a pH of 4.9), and solvent B (methanol). The gradient was 0 min, 0 % B; 3 min, 20 % B; 5.5 min, 20 % B; 11 min, 55 % B; 13.5 min, 95 % B; 16.5min, 95 % B; 17 min, 0 % B. The flow rate was 200 μL/min. Positive mode LC separation was achieved with a gradient of solvent A (0.025 % heptafluorobutyric acid, 0.1 % formic acid in H2O) and solvent B (acetonitrile) at 400 μl/min. The stationary phase was an Waters Atlantis T3, 3 μm, 2.1 mm × 150 mm column. The gradient was 0 min, 0 % B; 4 min, 30 % B; 6 min, 35 % B; 6.1 min, 100 % B; 7 min, 100 % B; 7.1 min, 0 % B. For both ionization modes, the injection volume was 10 μL and the QExactive Mass Spectrometer scanned in negative mode from *m/z* 70-1,000 at a resolving power of 70,000. For chiral analysis of serine, separations were performed with a Chirosil RCA (+) column (1.8 μm 4.6x150 mm). Separation was achieved with a gradient elution of solvent A (0.1% formic acid in water), and solvent B (0.1% formic acid in methanol). The gradient was 0 min, 0 % B; 3 min, 20 % B; 5.5 min, 95 % B; 6 min, 95 % B; 6.5 min, 0 % B; 8 min, 0 % B. The flow rate was 600 μL/min.

### Western Blots

Hemisected mouse brain tissue was lysed with tissue protein extraction reagent (TPER) (*Thermo Fisher Scientific*, 78501) and complete protease inhibitors (1 tablet/10 mL, *Roche*, 11836170001). Homogenized brain extracts were processed further via sonification using 5 seconds of pulsing and 5 seconds of rest for 2 rounds at 40 mA. The samples were then spun down at 14,000 rpms at 4°C in 25 minutes. Protein concentration was quantified with a bicinchoninic acid assay (BCA, *Thermo Fisher Scientific,* 23225). Lysates were separated by SDS-PAGE at 150V for 65 minutes on a 10% acrylamide gel and transferred to a PVDF membrane at 100V for 75 minutes. Membranes were blocked with 5% non-fat milk in 1X Tris-buffered saline with tween (TBST) for 1 hour at room temperature. Primary antibodies in 1x 5% non-fat milk were added to the membrane and probed overnight at 4°C. A secondary antibody in 1x 5% non-fat milk was added and kept on a rocker for 1 hour at room temperature. Membranes were then washed (3x) off once again in 1X TBST and imaged with Pierce enhanced chemiluminescence reagents (700 µL, Thermo Fisher Scientific, 32106) and exposed using Hyperfilm ECL film (*Cytivia*, 28906839). Primary antibodies used were as follows: anti-GLDC (rabbit, Thermo Fisher, PA5-22101, 1:800), anti-Vinculin (mouse, Millipore, V9131, 1:4000), anti-AMT (rabbit, Thermo Fisher, PA5-76454, 1:3000), anti-GCSH (rabbit, ProteinTech, 16726-1-AP, 1:1000), anti-phosphorylated PDH (Ser232) (rabbit, ProteinTech, 29582-1-AP, 1:6000), anti-PDH (rabbit, ProteinTech, 18068-1-AP, 1:6000), anti-DLAT (mouse, ProteinTech, 68303-1-Ig, 1:5000), anti- α Lipoic Acid (rabbit, Abcam, ab58724, 1:2000), and anti-SHMT2 (rabbit, ProteinTech, 11099-1-AP, 1:800).

Secondary antibodies were goat anti-mouse IgG (Bio-Rad, 1706516, 1:5000) and goat anti-rabbit IgG (Bio-Rad, 1706515, 1:5000).

### Statistical analysis

Statistical analysis was performed in GraphPad Prism (version 9.4.1) for both glycine analyses and western blots. The median levels ± SD was calculated for comparisons among treatment groups, and a two-way ANOVA with genotype and time as factors, followed by Tukey’s multiple comparisons test, was performed. A p-value of <0.05 was considered significant (*).

## Results

### Effects of *Gldc* mutation and animal age on brain glycine, serine, and responsiveness to folate deficiency

In prior studies we developed a computational mutation scale to predict mutations linearly proportional to disease severity, based on which we engineered mice homozygous for A394V mutation (predicted to be attenuated; *Gldc^ATT^*) as well as a second mouse carrying G561R, S562F (close to the active site) expected to be severe (*Gldc^SEV^*)^28^. Prediction of neurogenic severity was previously validated based on pre-natal disease, but post-natal studies were not undertaken. Importantly, absolute levels of post-natal brain glycine and serine remained unknown. To understand the effects of both mutation severity and animal age, we compared brains from a total of 65 mice containing attenuated and severe mutants as well as age and gender-matched heterozygous and wild type counterparts (**Fig. 1A**). For attenuated mutants we included two time points at ∼one and ∼ten months of age, but for severe mutants we used a single time point of ∼five months of age (due to difficulty in producing large numbers of severe mutants). As shown in **Fig. 1B**, we found that median glycine levels in brains of severe mutants were increased to 417.8 ng/mg brain tissue, while attenuated mutants at 1 and 11 months presented medians of 142.3 and 168.5 ng/mg, respectively. In wild type and heterozygous mice (across all ages), brain glycine levels ranged from 75-110 (with a median of 90 ng/mg). Together, these data suggested that in one month-old, attenuated mutants, brain glycine rose ∼1.5 fold, while severe mutants at five months, brain glycine was up by 4-fold and 2.5-fold compared to attenuated mutants at 1 and 11 months, respectively. Together, these data suggested that although animal age may play a role, the severity of the *Gldc* mutation primarily determined the extent of glycine elevation.

**Figure 1.**
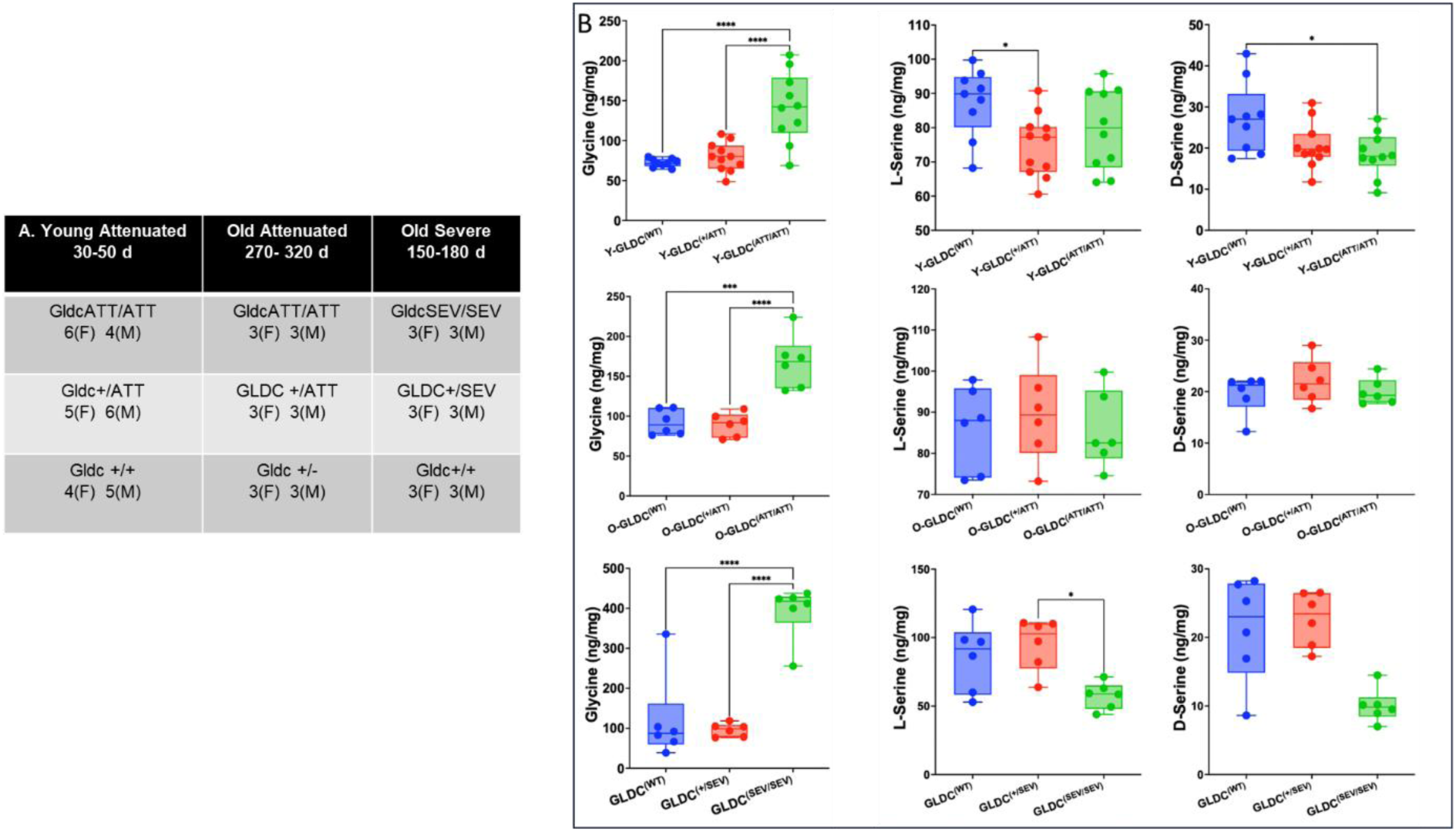
Age and genotype specific alterations in glycine and serine levels in brain tissue. **A.** Summary of experimental cohorts including young-attenuated (4-6 week), old-attenuated (9-11 month), and old severe (5-6 month) mice with indicated genotype and sex (M, males; F, females). **B.** LC-MS determination of L-serine and D-serine levels (in ng/mg) in brains of mutants (green) from young attenuated (Y-GLDC^ATT/ATT^), old-attenuated (O-GLDC^ATT/ATT^) and severe (GLDC^sev/sev^) mutants, their wild type (blue) and heterozygous (red) counterparts. Median glycine levels of Gldc^(SEV/SEV)^ mice was 417.8 ng/mg), that of Y-GLDC^ATT/ATT^ and O-GLDC^ATT/ATT^ mice were 142.3 and 168.5 ng/mg respectively, and wild type and heterozygous mice were at 90 ng/mg. Data are presented as individual values with medians ± SD. Significance was determined using one-way ANOVA.

Since serine and glycine can interconvert by the action of serine hydroxymethyl transferase 2 (SHMT2), we examined levels of L and D-serine in mouse brains. Compared to heterozygous counterparts, neither L or D serine were reduced in (young or old) attenuated mutants, but both were depleted in severe mutants (**Fig. 1C**). The reduction of serine in severe mutants suggested that SHMT2 does not run the reverse conversion of glycine to serine and L and D serine get converted into folate pathway (due to severe shortage of folates). We also found that severe mutants were reduced in betaine and increased in cystathionine and S-adenosylhomocysteine (SAH), while attenuated mutants only showed small increases in cystathionine (**Fig. 2** **A**). On the basis of these findings, we conclude that, although young and old attenuated mutants show increases in glycine and likely consume serine in cystathionine production, the amounts are low and do not result in marked change in pathways responsive to folate loss as shown by no changes in L or D serine, betaine, SAH. However, for severe mutants, there is a clear compensatory mechanism for the lack of 5- meTHF, and homocysteine is pushed toward cystathionine production rather than continuing the methionine cycle (which consumes a 5-meTHF unit), as proposed in the model in **Fig. 2B**.

**Figure 2.**
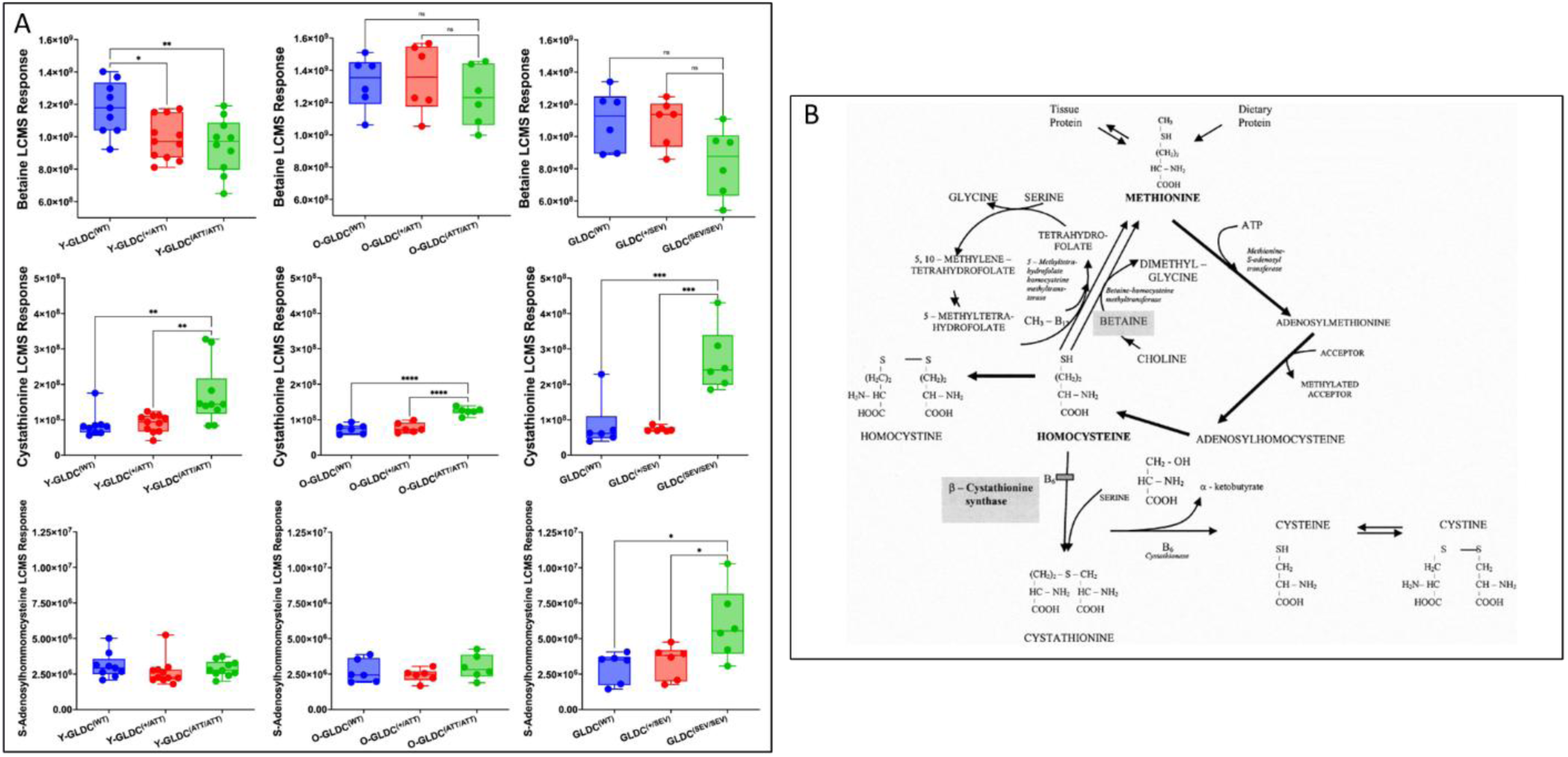
Age and genotype specific changes in betaine, cystathionine and S-adenosylhomocysteine (SAH) in brain tissue. **A** LC-MS response in young-attenuated Y- Gldc^(ATT/AAT)^ , old-attenuated O- Gldc^(ATT/AAT)^, severe Gldc^(SEV/SEV)^ mutants (green), their heterozygote (red) and wild type (blue) counterparts. **B.** Model summarizing severe and attenuated mutant responses. Severe mutants signal lack of 5-methyltetrahydrofolate and compensate by pushing elevated homocysteine toward cystathionine production, rather than continuing into the methionine cycle (which due reduction of betaine is blocked in production of 5-meTHF). Compared to heterozygotes, attenuated mutants do not show a reduction of betaine or increase in SAH. Serine in attenuated mutants may drive increases in cystathionine.

### Effects of *Gldc* mutation on the young-attenuated NKH brain proteome

We next undertook proteomic analyses to gain broader understanding of mutation-induced molecular changes in the brain. We examined the whole brain proteome of all 65 mice (**Fig. 3A**). Details of pre- processing of the proteomic data and statistical analyses are provided in Materials and Methods and Supplementary Table S1. Briefly, to identify proteins differentially expressed between disease and wild-type (WT) groups, we conducted age-matched two-sample t-tests for each protein, stratified analyses by phenotype and age group. Proteins with nominal p-values less than 0.01 were initially considered significant.

**Figure 3.**
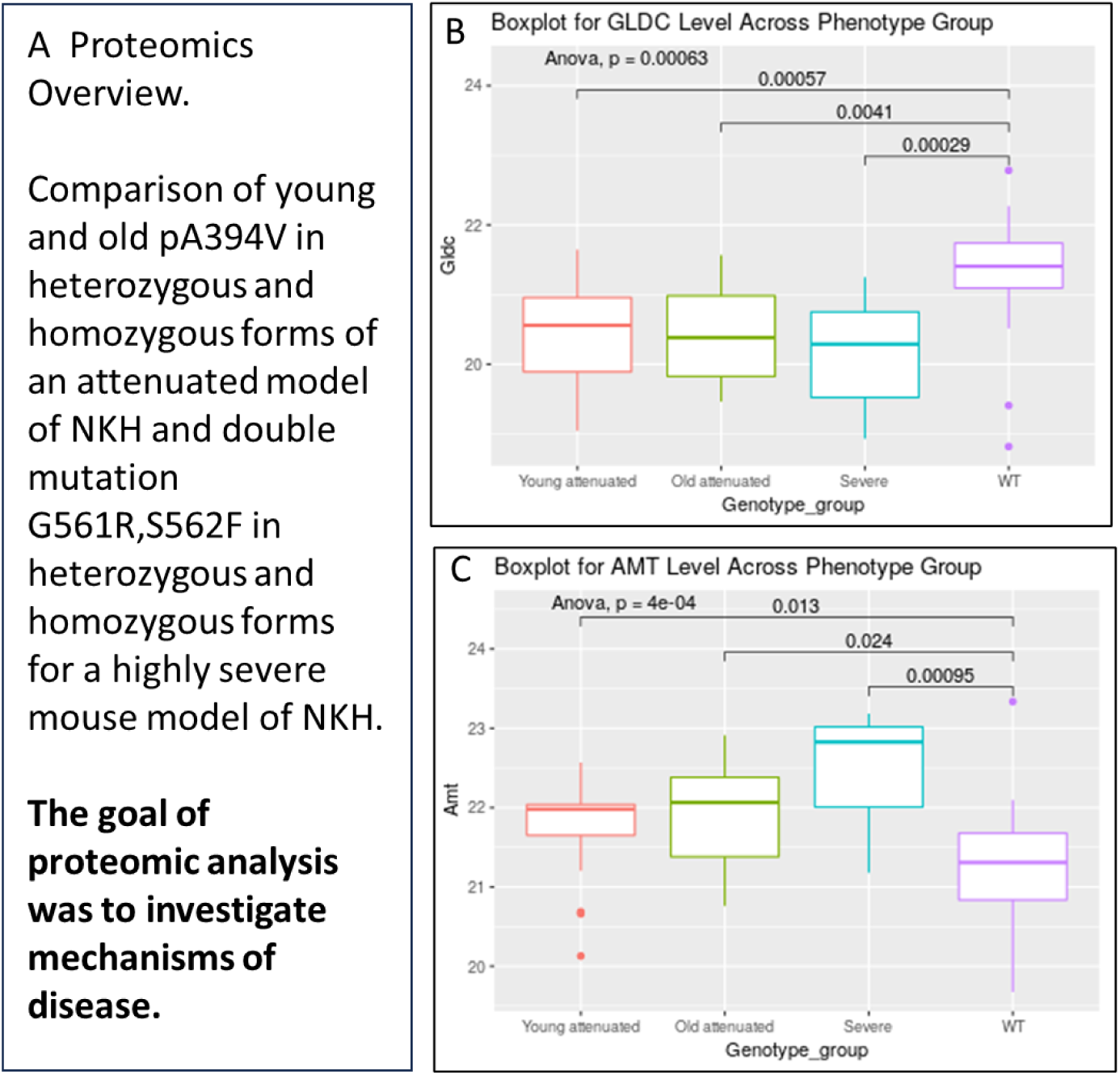
Proteomic analysis of differential expression patterns in mouse brain across genotype and phenotypes. **A.** Overview of experimental design. Brain tissue was analyzed and stratified by genotype and severity. **B.** Boxplots of GLDC protein levels across genotypes. **C.** Boxplot of AMT protein levels across genotypes. B-C boxplots represent interquartile ranges, with whiskers extending to the interquartile ranges and outliers shown as dots. Significance was determined using one-way ANOVA.

However, none of these remained significant after multiplicity correction using the Benjamini-Hochberg method.

To further explore the structure of the data, we applied unsupervised clustering techniques to the differentially expressed proteins. Across ∼6000 brain proteins, we investigated changes in levels of GLDC and a second GCS protein, aminomethyl transferase (AMT), since variants in these two genes combined cause 99% of NKH. We found GLDC levels were reduced in brains of attenuated (young and old) as well as severe mutant mice (**Fig. 3B**). In contrast, AMT was increased in mutants, particularly in severe mutants (**Fig. 3C**) consistent with much higher levels of compensatory responses for the lack of folates observed in severe mutant mice shown in **Fig. 2**.

Differential protein analyses of the whole brain proteome based on phenotype are shown in **Fig. 4**. These data suggested that brains of five-month-old severe mutants were better separated from their wild type and heterozygote mice, relative to attenuated counterparts at one month or 11 months. However, despite clear separations seen between metabolites across mouse groups (summarized in **Figs. 1-2**), we were unable to identify additional high value protein signatures shared across all three mutant phenotypes and that separated them from heterozygotes and wild type mice. One reason for this may be that we used only six mutants from older-attenuated and severe mutant groups, and these mice may present high levels of molecular heterogeneity intrinsic to differences in levels of *Gldc* deficiency. Notably, patients also show high levels of clinical heterogeneity with presentation of over 50 symptoms and no single symptom shared by all patients^27^. Nonetheless, reduction of GLDC protein across all three proteomes, provided molecular basis for glycine elevation in brains of all attenuated and severe mutants (shown in **Fig. 1**). The sharp elevation seen in AMT in the severe mutant proteome is also consistent with data presented in **Figs 1-2**, predicting that these mouse brains present the greatest responsiveness to folate loss.

**Figure 4.**
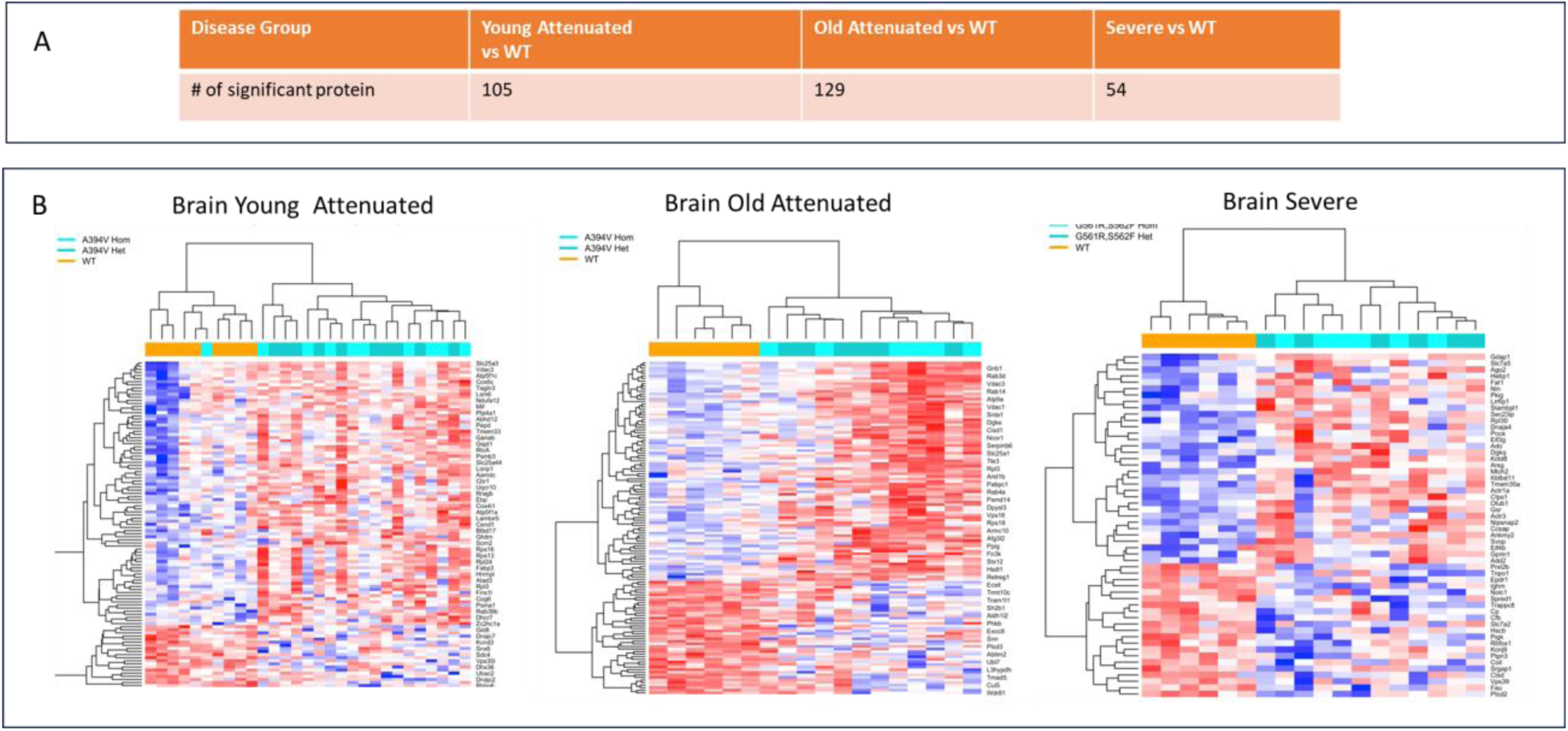
Differential protein expression andunsupervised clustering based on phenotype. Protein expression levels between age matched diseased group and WT group was investigated using t-test. **A.** Number of differentially expressed proteins in each phenotype-age group (p < 0.01). **B.** Unsupervised clustering of the differentially expressed proteins for groups described in A. The homozygous mouse that clustered with wild type was confirmed by genotyping, suggesting young attenuated mutants show high degree of phenotypic heterogeneity. Graphs use mean imputed data for missing values; t-test p< 0.01. Adjusted p values were not significant.

### Effects of Gldc mutation on GCS and the PDH complex proteins

GLDC catalyzes the first and the rate limiting step of glycine cleavage in the GCS. It forms a complex with GCSH protein, needed for the transfer of the aminomethyl group released from glycine to the AMT protein. Dihydrolipoamide dehydrogenase (DLD), the fourth protein of the GCS, resets the glycine cleavage reaction by re-oxidizing the lipoyl moiety of GCSH. Since GLDC has direct molecular interactions with GCSH^27^, a reduction in the former may antagonize the stability of the latter. GCSH is a lipoyl transfer protein of the GCS, but it also mediates lipoylation of other mitochondrial proteins^37^ , including components of the PDH complex^38^ , which play key roles in mechanisms of energy metabolism in the brain.

To identify early mechanisms of mitochondrial energy mechanisms engaged by defect in GLDC, we focused on attenuated mutants aged one month. As shown in **Fig. 5A-D**, whole brains of these mice showed greater than 80% reduction of GLDC, GCSH decreased by 25% while AMT increased by 15%. Together, these data strongly suggested that GLDC was quantitatively reduced throughout the brain. Moreover, since over 90% of GLDC in the mouse brain is present in astrocytes^33^, depletion of GLDC principally occurs and affects other GCS effectors in astrocytes. However, unlike GLDC, GCSH functions in all cells of the brain and hence its 25% reduction in whole brain lysates, underestimates its real decrease in astrocytes. The concomitant decrease of GLDC and GCSH implicated potential broader effects on mitochondrial energy mechanisms of astrocytes.

**Figure 5.**
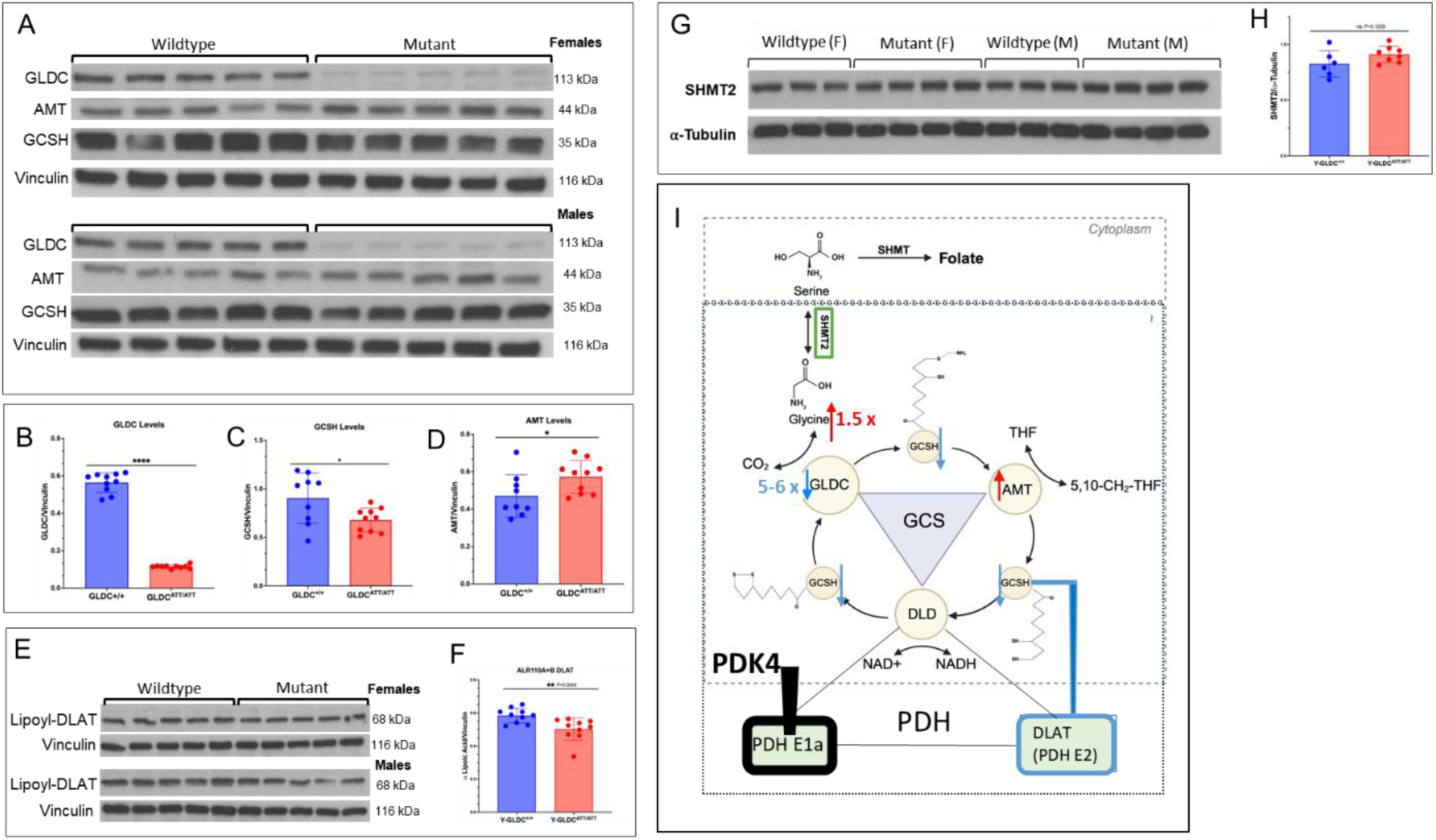
pA394V mutation reduces levels of GLDC, GCSH and lipoylation of DLAT in astrocytes. **A.** Representative western blots showing GLDC, GCSH and AMT in whole brain tissue from young-attenuated mutants compared to and wild type for female and male mice. **B-D.** Corresponding quantification of GLDC (reduced 5-6-fold), GCSH (reduced 1.5-fold) and AMT in wild type and mutant brains. **E-F** Representative western blots showing lipoylated DLAT in brains of young-attenuated mutants compared to and wild type for female and male mice and associated quantification. **G-H.** Representative western blots showing SHMT2 in brains of young-attenuated mutants compared to and wild type for female and male mice and associated quantification. **I**. Astrocyte model summarizing changes seen in the GCS and PDH complexes. Blue indicates reduction in GLDC, GCSH and lipoylation of DLAT. AMT is raised (red) and SHMT2 is unchanged. Astrocyte PDK4 keeps PDHE1α unchanged in a highly phosphorylated, inactivated state (black), suggesting reduction in lipoylation of DLAT may trigger alternate mitochondrial mechanisms of energy metabolism.

Since GCSH is a lipoyl-transfer protein, we considered whether the reduction in its levels may affect lipoylation of other mitochondrial proteins. We examined the effect on lipoylation of dihydrolipoamide S-acetyl transferase (DLAT), which forms the E2 subunit of the PDH complex (with DLD being the E3 subunit). As shown in **Fig. 5E-F**, lipoylation of DLAT was found to be measurably decreased (by 13%) in mutant compared to wild type brain. Again, since this reduction occurs due to mutation in GLDC in astrocytes, we expect the greatest decline of DLAT-lipoylation to also be in astrocytes (and 13% reduction seen in whole brain lysates is likely to underestimate reduction in astrocytes). Levels of AMT were increased by 15% in the young- attenuated mutants and those of SHMT2 were unchanged (**Fig. 5G-H**), strongly suggesting that AMT and

SHMT2 levels were not impaired, and the observed depletion of GCSH and DLAT-lipoylation were due reduction in GLDC. Thus, as summarized in the model in **Fig. 5I** depletion of GLDC (rather than glycine elevation or serine reduction) lowers GCSH and lipoylation of DLAT with potential to impact mitochondrial energy mechanisms of astrocytes.

### Effects of *Gldc* mutation on astrocyte signatures of β−oxidation of fatty acids, glycolytic intermediates, and activation of neuronal PDH

Since DLAT is a key component of PDH, reduction in its lipoylation is expected to activate mitochondrial mechanisms of energy metabolism. However, as shown in the model in **Fig. 5I**, astrocytes do not favor activation of their PDH, since this may antagonize neurometabolic support that astrocytes provide to neurons^39^. To identify alternate mitochondrial mechanisms, we undertook Ingenuity Pathway Analyses (IPA) of the attenuated mutant proteome (p < 0.05; Supplemental Table S2). As shown in **Fig. 6A**, we found that two of the top 20 altered pathways were mitochondrial pathways of oxidative phosphorylation and fatty acid β-oxidation. Oxidative phosphorylation is seen in both neurons and astrocytes, but the former prefer glucose as an energy source, while β-oxidation of short chain fatty acids is characteristic of activation of astrocyte mitochondria^16^.

**Figure 6.**
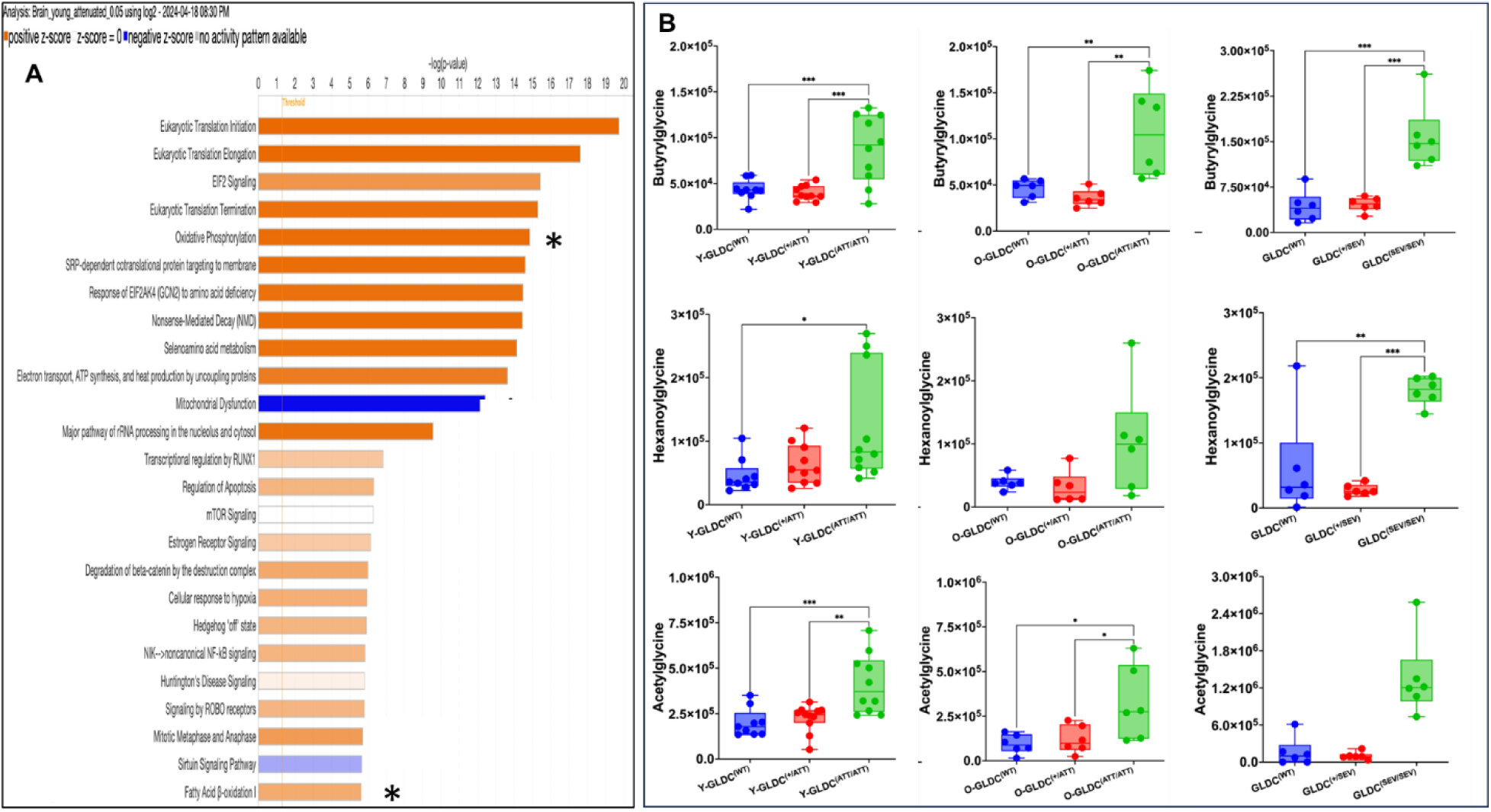
Age and genotype specific changes in mitochondrial pathways of energy metabolism. **A.** Ingenuity Pathway Analyses (IPA) predicts the proteome of young Gldc^(ATT/ATT)^ mice is enriched in two mitochondrial pathways of energy metabolism, namely oxidative phosphorylation and fatty acid beta-oxidation activity. Pathways with significant enrichment are ranked by –log(p-value), where orange bars indicate upregulated pathways, blue bars indicate downregulated pathways, and gray indicates no directional change. **B.** LC-MS response in changes in acylglycines (namely butyrylglycine and hexanoylglycine) as well as acetylglycine in young-attenuated (Y- Gldc^(ATT/AAT)^), old-attenuated (O-Gldc^(ATT/AAT)^) and severe (O-Gldc^(SEV/SEV)^) mutants (green), and their heterozygous (red) and wild type (blue) counterparts, suggesting mutants activate β-oxidation of short chain fatty acids, an astrocyte mechanism of mitochondrial energy metabolism.

We therefore screened brains for signatures of production of short chain fatty acid acids. As shown in **Fig. 6B**, we followed hexanoylglycine, butyrylglycine and acetylglycine as sensitive signatures of glycine capture by newly produced intermediates of β- oxidation of short-chain fatty acids and activated acetyl groups that could be easily differentiated between mutant and wild type outputs. In young-attenuated mutants, the levels of butyrylglycine and acetylglycine were significantly elevated compared to both wild type and heterozygotes. Increase in hexanoylglycine was statistically significant only relative to wild type. Analyses of old attenuated mutants and severe mutants showed additional increases in all three acylglycines and acetylglycine, with greatest changes seen in severe mutants (**Fig. 6B**). This suggested that fatty acid β- oxidation is stimulated as an early mitochondrial response in young- attenuated mutants that become elevated as mice age or in response to increase in the severity of the mutation.

We also examined glycolytic breakdown products of fructose-1,6-bisphosphate (F16P), 2,3- bisphosphoglycerate (2,3-BPG) and phosphoenolpyruvate (PEP) that occur in the cytoplasm as precursors to pyruvate (before it is imported into mitochondria). As shown in **Fig. 7**, we found glycolytic metabolites were sharply raised in severe mutants at 5 months. They increased to a lesser degree in old-attenuated mutants at 11 months. But notably, young-attenuated mutants showed no increase in glycolytic intermediates. A stimulant of mitochondrial energy mechanisms guanidino acetic acid (GAA) was significantly increased across attenuated and severe mutants (with highest levels associated with severe). A second stimulant, 5- aminoimidazole-4-carboxamide-1-β-D-ribofuranoside (AICAR) which signals starvation and acts as an activator of AMP-activate protein kinase (AMPK) was significantly raised in severe mutants compared to heterozygotes and wild type mice, but not in attenuated mouse brains.

**Figure 7.**
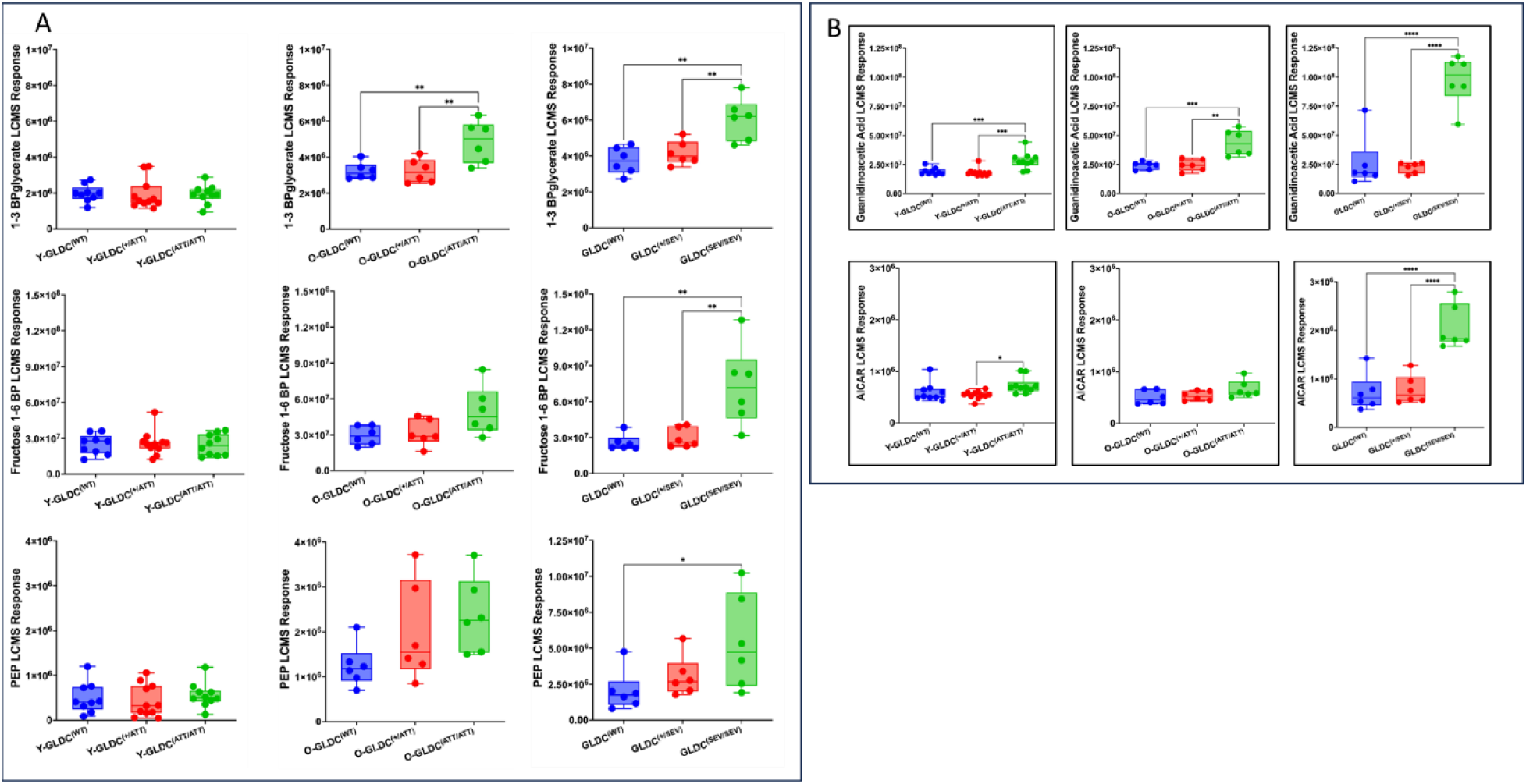
Age and genotype specific changes in glycolytic intermediates and other factors of energy metabolism. **A.** LC-MS responses in changes in glycolytic intermediates in young-attenuated (Y-Gldc^(ATT/AAT)^) old-attenuated (O-Gldc^(ATT/AAT)^) and severe (O-Gldc^(SEV/SEV)^) mutants (green), and their heterozygous (red) and wild type (blue) counterparts. **B**. LC-MS responses in changes in GAA and AICAR in young-attenuated (Y-Gldc^(ATT/AAT)^) old-attenuated (O-Gldc^(ATT/AAT)^) and severe (O-Gldc^(SEV/SEV)^) mutants (green), and their heterozygous (red) and wild type (blue) counterparts.

Together, the data in **Fig. 6-7** suggested that a primary effect of *Gldc* mutation was stimulation of mitochondrial β-oxidation of short chain fatty acids and activated acetyl groups in astrocytes^16^. Increased glycolytic intermediates in older attenuated mutants and severe mutants is consistent with PDH dysfunction via decreased lipoylation. Elevation of GAA rather than AICAR suggests that GAA, a primary stimulant underlying the observed increase in signatures of mitochondrial oxidation of fatty acids in young-attenuated mutants. That AICAR is not raised, suggests while the young-attenuated mutant astrocytes activate alternate mechanisms of energy metabolism they are not yet starved, but in early stages of energy depletion.

Unlike the situation in astrocytes, in neurons phosphorylation of the E1 subunit of PDH (PDHE1α) is inversely associated with their mitochondrial energetics ^40^. We therefore compared ratios of its phosphorylated and unphosphorylated forms in brains of mutant and wild type mice. As shown in **Fig. 8A**, mutant brains showed significantly lower levels of phosphorylation, suggesting significant activation of PDH-mediated mitochondrial metabolism of neurons. However, since neurons largely lack GLDC^33^ the data suggest activation of neuronal PDH arises due to defect in astrocyte GLDC underlies, possibly through astrocyte-neuronal communication such as via the astrocyte-neuron lactate shuttle (ASNLS) shown in **Fig. 8B**. The ASNLS is a major mechanism of metabolic communication between astrocytes and neurons. It enables lactate (derived from pyruvate) in astrocytes to be delivered to neurons, where it is converted back to pyruvate to boost PDH-dependent mitochondrial energy metabolism^2^. Thus, in summary, the model in **Fig. 8B** states that quantitative reduction of GLDC, GCSH and DLAT lipoylation in astrocytes coupled with PDK4-mediated heavy phosphorylation of PDHE1α which limits pyruvate consumption via decarboxylation^41^ , triggers alternate mechanisms of mitochondrial activation such as β-oxidation of fatty acids. Moreover, astrocyte conversion of increased pyruvate to lactate, which is transported via the ANLS to neurons, drives PDH activation.

**Figure 8.**
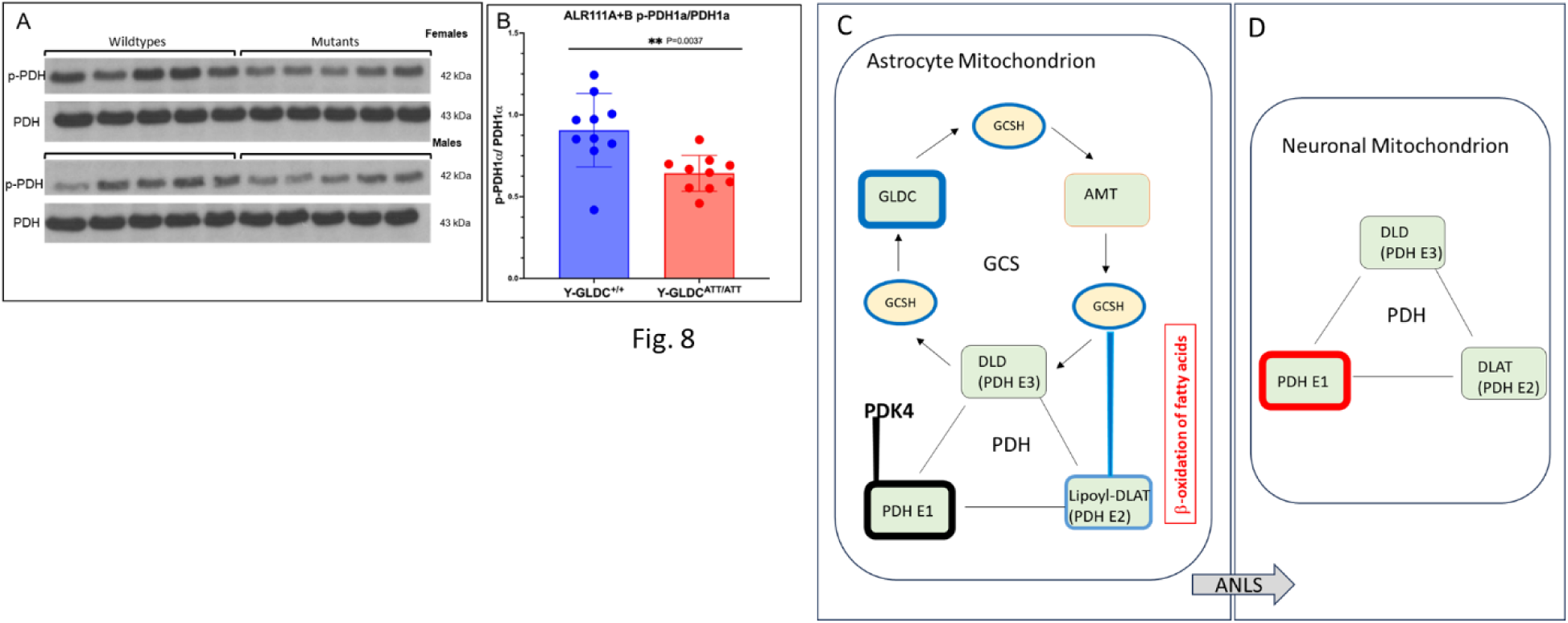
Deficiency in GLDC decreases PDH phosphorylation of neurons. **A.** Representative western blots showing levels of phospho-PDHE1a and total PDHE1a levels in whole brain tissue from young-attenuated mutants (Y-GLDC ^ATT/ATT^) and wild type for female and male mice. **B.** Corresponding quantification of the ratio of p-PDHE1a/PDHE1a in wild type and mutants. **C-D** Model summarizing changes seen in the GCS and PDH complexes of astrocyte and neurons. **C. Astrocytes**. Blue indicates reduction in GLDC, GCSH and lipoylation of DLAT. AMT is raised (red) and SHMT2 is unchanged (not shown; see Fig. 5). Astrocyte PDK4 keeps PDHE1a unchanged (black) in a highly phosphorylated state. Beta-oxidation of short chain fatty acids is activated (red). **D. Neurons**. Activation of PDH (red) detected in response to defect in astrocyte GLDC, suggests communication via the astrocyte-neuron lactate shuttle (ANLS).

## Discussion

Our studies show that in our NKH mouse models, post-natal changes in brain glycine, D and L serine, betaine, cystathionine, GAA, and SAH are proportional to the severity of the mutation defect in GLDC. Unexpectedly, in young-attenuated mouse brains, the direct enzymatic consequence was limited to 1.5-fold elevation in glycine. There were no metabolite changes reflecting responsiveness to folate depletion or alterations in the levels of SHMT2. But these brains showed greater than five-fold reduction in GLDC levels. This suggested that at reduced protein levels, the catalytic turnover of GLDC may be increased to limit metabolite changes. Further, the massive reduction of GLDC protein levels may inflict non-enzymatic properties which have never previously been reported for either NKH or GLDC. However, the wide spectrum of disease symptoms associated with NKH is consistent for both enzymatic and non-enzymatic properties of mutant GLDC (and other GCS components) to play deleterious roles in the brain.

The non-enzymatic consequences of GLDC depletion evident in our mouse model were reflected by reduction in both GCSH and lipoylation of DLAT, a component of PDH. Defect in GCSH is known to cause NKH and lipoate deficiency^42^. But when disease is initiated by defects in GLDC, the changes are expected to be in astrocytes since they are the major cell type where GLDC localizes in the brain. This coupled with the fact that astrocytes make up the majority of cells in the brain^43^ enabled use of our NKH mutant mice as a model system to understand astrocyte functions in the NKH brain. Decrease in lipoylation of DLAT with concomitant increase in signatures of β−oxidation of fatty acids confirmed that defect in GLDC activated alternate energy mechanism of astrocyte mitochondria. That these changes are seen in whole brain lysates suggest that they are widespread in astrocytes throughout the brain. Moreover, these astrocyte changes were seen in young-attenuated mice, suggesting they occur early, from the onset of disease in the mouse and persist throughout post-natal development.

Prior studies have suggested that in lung cancer cells, pyruvate metabolism decreases on GLDC inhibition^44^ In hepatocellular carcinomas elevation of GLDC increased lipoylation of DLAT to promote tumorigenesis^45^ and knock down in GLDC decreased DLAT lipoylation to suppress PDH activation and tumor growth. However, in astrocytes PDK4 ensures that PDH remains highly phosphorylated and blocked in pyruvate decarboxylase activity and this singular characteristic prevents astrocyte PDH activation. But we detected neuronal activation of PDH because of mutation of GLDC in astrocytes. In the brain, a major function of astrocytes is to support neurons particularly with metabolic homeostasis. The transport of lactate from astrocytes to neurons, via the ANLS is an important hypothesis of metabolic cooperation between astrocytes and neurons^46^. In astrocytes PDK4 keeps PDH highly phosphorylated and blocked in pyruvate decarboxylase activity and this unique feature promotes pyruvate conversion to lactate and fuel the ANLS^47^. Our finding that Gldc mutation-induced defect in DLAT lipoylation in the PDH complex of astrocytes occurs with concomitant stimulation of neuronal PDH, suggests a role for the ANLS in communicating how defect in GLDC in astrocytes may alter mitochondrial PDH activity in neurons (**Fig. 8B**).

The attenuated mutant pA394V used in our studies is the murine orthologue of human pA389V, a prevalent variant with reported expansion in patient populations^48^ with associated symptoms of poor feeding, hypotonia and seizures ^48^. We have recently reported that as these mutants mature from one to five months, they develop long term neurological disease and death, which is associated with decline in astrocytes (but not neurons, oligodendrocytes, or microglia), consistent that the defect in GLDC exerts a primary antagonistic effect on astrocytes. Why activation of mitochondrial energy mechanisms cause astrocyte loss is not known, but one possibility is that fatty acid oxidation-induced downstream stimulation of oxidative phosphorylation generates reactive oxygen species (ROS)^16^ that is detrimental to astrocytes. Ketogenic diets shown to benefit severe and attenuated NKH patients^49^ ^50^ have been ascribed to reduction of glycine, but they are known to improve brain energy metabolism^51^ which may contribute to their success in treating NKH. By enabling neurons to use ketones, ketogenic diets may help reduce their dependence on oversupply of ANLS as well as tamp-back activation of β-oxidation of fatty acids caused by mutation in astrocyte GLDC and thereby mitigate mitochondrial ROS in astrocytes. But the availability of ketones to fuel neuronal PDH activation may increase oxidative phosphorylation and ROS in neurons and thereby may limit the value of ketone diets to mitigate long term disease. We conclude by noting that our studies provide the first evidence that GLDC directly plays a role in regulating mechanisms of mitochondrial energy metabolism in the brain. Our attenuated and severe NKH mutant mouse models provide tools to further delineate fundamental mechanisms of energy regulation in the brain.

## Acknowledgments

We thank the members of the Haldar lab for helpful discussions during the course of the work.

## Animal facility

We thank the Freimann Life Sciences Center (FLSC) and staff at the University of Notre Dame, for providing excellent services for mouse housing, caring, breeding, blood drawing, overall monitoring and veterinarian support.

## Regulatory Approval

All procedures were approved by the Institutional Biosafety Committee (IBC), University of Notre Dame. Design of animal studies and procedures regarding their use were approved by the Institutional Animal Care and Use Committee (IACUC) University of Notre Dame.

## Author Contribution

Alejandro Lopez-Ramirez – conceptual design, experimentation, analyses and interpretation of studies in Fig. 1, 2, 4, 5, 6, 7 and 8, colony management organ harvest, visualization of results, writing of related methods and figure legends, abstract, introduction, results, discussion and overall primary drafting and editing of the manuscript.

Andrew Worth - conceptual design, experimentation, metabolomic analyses and interpretation of studies in Fig. 1, 2, 3 4, 6 and 7, visualization of results, writing of related methods and editing of the manuscript.

Lindsay Wang – proteome data processing and cleanup, along with analysis and data Fig 4. Generation.

SalihaYilmaz – oversight of bioinformatic analysis, proteome data processing and Fig 4 generation, interpretation of studies and editing of the manuscript.

Sebastian Hayes – Oversight and generation of proteomic data

Joseph Farris – conceptual design, analyses and interpretation of studies in Fig. 1, 2, 4, 5, 6 and 7 and editing of the manuscript.

Md Suhail Alam – conceptual design and experimentation (Figure 1), colony management organ harvest and editing the manuscript.

Prasad Padmanabhan – colony management, organ harvest and editing the manuscript.

Kasturi Haldar – overall conceptualization, design and supervision of entire project, design, all data analyses and visualization of results, primary drafting and editing of manuscript, funding acquisition.

